# A genome-wide CRISPR functional survey of the human phagocytosis molecular machinery

**DOI:** 10.1101/2022.09.08.507072

**Authors:** Patrick Essletzbichler, Vitaly Sedlyarov, Fabian Frommelt, Didier Soulat, Leonhard X Heinz, Adrijana Stefanovic, Benedikt Neumayer, Giulio Superti-Furga

## Abstract

Phagocytosis, the process of engulfing large particles by cells, is a multilayered biological activity driving tissue clearance and host defense. Dysregulation of phagocytosis is connected to autoimmunity, accumulation of toxic disease proteins, and increased risks for infections. Despite its importance and multiple roles, we lack a full understanding of the cellular machinery involved in executing and regulating the process, including the coordination with other cellular events. To create a functional map in human cells, we performed a reporter- and FACS-based genome-wide CRISPR/Cas9 knock-out screen that identified 716 genes. Mapping the gene hits to a comprehensive protein-protein interaction network annotated for functional cellular processes, allowed to highlight those protein complexes identified multiple times, to identify missing components of the cellular phagocytosis network, and to suggest functional partition among complexes. We validate complexes known to be involved, such as the Arp2/3 complex, the vacuolar-ATPase-Rag machinery, and the Wave-2 complex, as well as processes previously not or only poorly associated with phagocytosis. Among the novel, phagocytosis-relevant cellular functions validated are the oligosaccharyltransferase complex (MAGT1/SLC58A1, DDOST, STT3B, and RPN2) as well as the hypusine pathway (eIF5A, DHPS, and DOHH). Overall, our network of phagocytosis regulators and effectors maps elements of cargo uptake, cargo shuffling and cargo biotransformation through the cell, providing a valuable resource for the identification of potential novel drivers for diseases of the endo-lysosomal system. We further propose that our approach of mining and integrating publicly available protein-protein interaction data with datasets derived from reporter-based genome-wide screens offers a broadly applicable way to functionally map biological processes onto the molecular machinery of the cell.

**Summary blurb:** The validation and interpretation of a FACS reporter-based genome-wide CRISPR/Cas9 knock-out screen through protein-protein interaction data yields a comprehensive view of the molecular network regulating and executing phagocytosis in human cells.

## Introduction

Phagocytosis is a multifaceted, integrated biological process, that involves the engulfment of large (≥0.5 µm) particles under strict coordination over time and space, with an immunological and homeostatic purpose (Flannagan et al., 2012; Gordon, 2016; Underhill and Goodridge, 2012). It is a key defense and neutralization mechanism against pathogens and at the same time involved in the clearance of apoptotic cells (Arandjelovic and Ravichandran, 2015), in the processes of wound healing (Giulian et al., 1989), in tumor cell phagocytosis (Feng et al., 2019; Kamber et al., 2021), in microglial phagocytosis (Podleśny-Drabiniok et al., 2020), and in the resolution of tissue injury (Gerlach et al., 2021). Among other phenomena, dysregulated phagocytosis or aberrant phagosome maturation can promote Alzheimer’s disease, allow pathogens like bacteria or viruses and cancer cells to escape destruction, or promote autoimmunity through incomplete degradation of cell debris and DNA (Arandjelovic and Ravichandran, 2015; Podleśny-Drabiniok et al., 2020).

Material taken up by a cell is contained in a membrane-bound vacuole called phagosome that matures through a series of highly organized membrane fusion and fission events, altering its composition and gradually acidifying its pH (Flannagan et al., 2012). A freshly formed phagosome (pH ∼7.4) fuses first with early endosome acquiring among others the GTPase Rab5. Then it undergoes a series of maturation steps, yielding the late phagosome marked by the exchange of Rab5 to Rab7 and the acquisition of additional vacuolar ATPase proton pumps, which acidify the phagosome to pH 5.5-6.0, forming the ideal degradative environment (Flannagan et al., 2012).

Systematic searches for phagocytosis regulators have been widely undertaken with RNAi-based screens in cultured Drosophila cell models (Kocks et al., 2005; Philips et al., 2005; Rämet et al., 2002) Until recently, such genetic screens have been lacking for mammalian cell systems (Haney et al., 2018; Lindner et al., 2021; Sedlyarov et al., 2018; Yeung et al., 2019). The advancement of gene-editing tools, in particular the discovery of CRISPR/Cas9 systems (Cong et al., 2013; Jinek et al., 2013; Mali et al., 2013), and increasingly sophisticated read-out systems, now allows conducting large genome-wide knock-out screens in mammalian cells at high precision (Cong et al., 2013; Sanjana et al., 2014; Shalem et al., 2015).

Once it is established that a reporter system faithfully represents the biology under investigation and once validation and calibration of the system allow it to display the necessary signal-to-noise ratio, pooled fluorescent-activated cell sorting (FACS)-based genetic screens offer a powerful method to genetically map a large variety of biological processes (Bock et al., 2022; Doench, 2018).

Here we present a FACS-based genome-wide CRISPR/Cas9 knock-out screen for phagocytosis using a reporter assay (Colas, 2020) that we previously employed for focused genetic screening (Sedlyarov et al., 2018) as well as for measuring the maturation of the phagolysosome (Heinz et al., 2020). As in vitro macrophage cell model we chose phorbol myristate acetate (PMA)-differentiated human THP-1 cells, a classical cell model to study phagocytosis and immune modulation (Chanput et al., 2014; Fleit and Kobasiuk, 1991).

The study reported here represents a comprehensive genetic assessment of all the non-essential cellular functionalities involved in phagocytosis, including its regulation, dynamic phagosomal maturation processes, and movement by the complex cell cargo transport machinery. Additionally, the study highlights the power of combining the information from functional genetic screening and publicly available protein-protein interaction data, representing the physical base for concerted cellular activities. We present the efficient guiding of validation as well as the rationalization of the genetic results in light of the annotated cellular machinery and its individual components.

## Results

### A FACS reporter-based screen for modulators of phagocytosis

Phagocytosis proceeds through defined key steps, including the uptake of cargo, the systematic coordinated transport to the lysosome, and proper acidification. We set out to identify genes potentially involved in the entire process by making use of pooled CRISPR/Cas9 knock-out screening in combination with reporter-based flow cytometry assisted cell sorting. For this, we chose the myeloid cell line THP-1 (Chanput et al., 2014) and a PMA-based differentiation protocol that induces in these cells a macrophage-like phenotype (Fleit and Kobasiuk, 1991). To validate the experimental setup we tested actin polymerization using Cytochalasin D (Carter, 1967) and lysosome-dependent acidification using Bafilomycin A1 (**Figure 1b-c**) (Dröse et al., 1993). Next, we introduced a genome-wide CRISPR/Cas9 knock-out library, targeting all protein-coding human genes with 70,948 sgRNAs as well as 142 control sgRNAs (Hart et al., 2017). We use the lentiCRISPRv2 single vector driving the expression of a sgRNA-cassette as well as Cas9 developed by Sanjana and colleagues (Sanjana et al., 2014).

**Figure 1:**
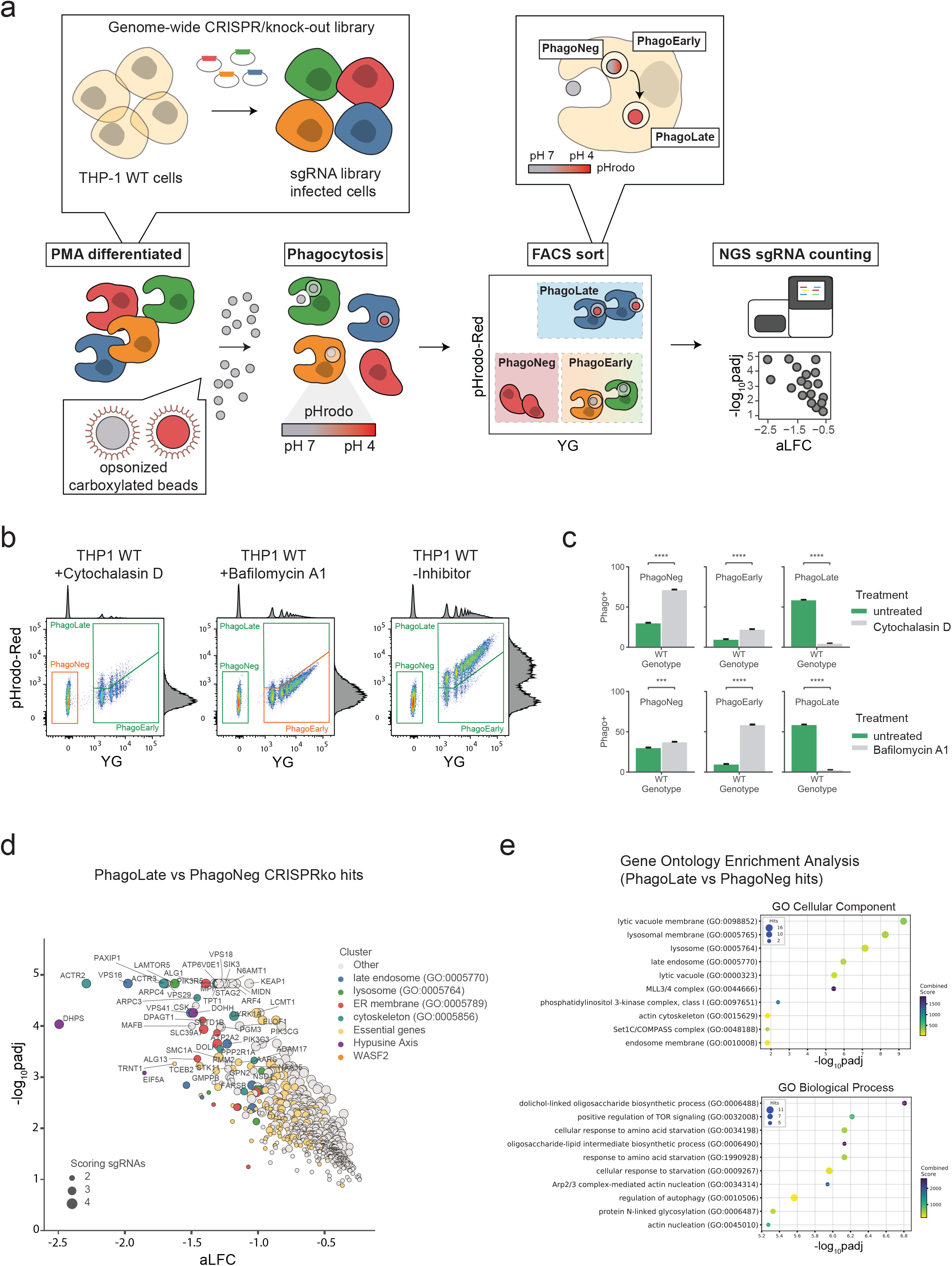
**a**. Schematic overview of the FACS-based genome-wide CRISPR/Cas9 knock-out screen to identify modulators of phagocytosis. **b**. Representative flow cytometry scatterplots of phagocytosis assays. PMA-differentiated THP-1 cells were either treated with the control drugs Cytochalasin D or Bafilomycin A1 or were left untreated and then incubated with opsonized, dual-colored reporter beads. Each dot corresponds to one measured cell: the intensity of the pH-insensitive dye is represented on the x-axis (YG), and the signal of the pH-sensitive dye, which increases signal with decreased pH is displayed on the y-axis (pHrodo-Red). Double-negative cells were sorted as phagocytosis-negative (PhagoNeg), single-positive cells (YG high signal but low pHrodo-Red signal) were grouped as early stages of phagocytosis and double-positive cells (YG high siganl and high pHrodo-Red signal) were classified as cells that underwent phagocytosis and phagosome acidification (PhagoLate). Cells treated with the control drug Cytochalasin D remained mostly in the PhagoNeg gate and cells treated with Bafilomycin A1 stayed in the PhagoEarly gate. The marginal intensity distributions are illustrated on the side of the plot. **c**. Barplot showing the quantification of the different fractions of cells gated in (b). Data are mean ± 95% confidence interval from 3 replicates. ***p ≤ 0.001, ****p ≤ 0.0001; by Welsh’s t test. **d**. Volcano plot showing the 716 genes depleted in the PhagoLate population versus the PhagoNeg population. The x-axis shows the average log_2_ -fold change (aLFC) calculated for all sgRNAs per gene against the y-axis, representing the statistical significance as -log_10_(p_adj_). The size of the dot indicates the amount of sgRNAs changing significantly for the particular gene. **e**. Gene Ontology (GO) enrichment analysis (two-sided Fisher’s exact test, p value adjusted for multiple testing) for genes depleted in the PhagoLate population seen in (d) for GO Cellular Components and GO Biological Processes. The x-axis shows the significance of enrichment (-log_10_-transformed p value adjusted for multiple testing) against the y-axis showing the top 10 enriched terms, sorted by adjusted p value.

To measure phagocytic uptake of cargo as well as the transport and acidification of the resulting phagolysosome, we employed a flow cytometry readout based on opsonized 1.75 µm latex beads coated with the pH-sensitive dye pHrodo (Colas et al., 2014) and successfully adopted for genetic screening (Sedlyarov et al., 2018). While cells that were fully functional in phagocytosis and acidification appeared in the PhagoLate fraction, cells still capable of phagocytosis, but deficient in acidification, emerged in the PhagoEarly fraction (**Figure 1a**). In contrast, cells that were completely deficient in phagocytosis appeared in the PhagoNeg fraction. Since the cytoskeleton has a key role in the formation of the phagocytic cup, as well as for the uptake of material, we employed the cytoskeletal inhibitor Cytochalasin D to show the uptake of the reporter would be blocked efficiently. Cytochalasin D treated cells emerged predominantly in the PhagoNeg fraction, indicating that their phagocytic defect stemmed from an uptake issue (**Figure 1b-c**). We treated the cells with a vacuolar-ATPase inhibitor Bafilomycin A1, to show that acidification of the lysosome was required for cells to emerge in the PhagoLate gate. The uptake of the reporter itself remained unaffected, which led to a rise of cells in the PhagoEarly gate (**Figure 1b-c**).

### A genome-wide screen identifies known and novel modulators of phagocytosis

After setting up the phenotypic assay to measure phagocytosis in THP-1 cells, we next performed a genome-wide screen for modulators of phagocytosis in quadruplicates following the workflow portrayed in **Figure 1a**. We introduced first a genome-wide CRISPR knock-out library into THP-1 cells and selected cells with puromycin before we induced a macrophage-like phenotype via PMA differentiation. The period chosen for further cell incubation was optimized to yield approximately one-third of cells in the PhagoNeg fraction and two-thirds of the cells in the phagocytosis- and acidification-positive fractions (PhagoEarly and PhagoLate; 3 hours). Cell fractions were gated, FACS sorted and processed separately allowing for comparison of sgRNA abundance within each population and replicate (**Suppl Fig 1a-b**).

By comparing sgRNAs that were depleted in PhagoLate compared to PhagoNeg we identified in total 716 genes to be involved either in phagocytosis itself or in phagosome acidification (**Figure 1d and Suppl Table 1**). Enrichment analysis for “Cellular components” GO terms yielded among others as strongly enriched compartments the lytic vacuole membrane, lysosome, and late endosome (adj. p-value < 1.08 × 10^−6^). These components are all part of cellular processes and pathways which are well known to be involved in phagocytosis (**Figure 1e**). Enriched biological processes included the dolichol-linked oligosaccharide biosynthetic process, TOR signaling, starvation response as well as actin nucleation (adjusted p-value <5.30 × 10^−6^) (**Figure 1e**). RNA-sequencing of THP-1 cells treated with the reporter was further used to confirm that genes identified as phagocytic regulators were expressed in THP-1 cells across a wide range of expression (**Suppl Fig 2a-b** and **Suppl Table 3**).

**Figure 2:**
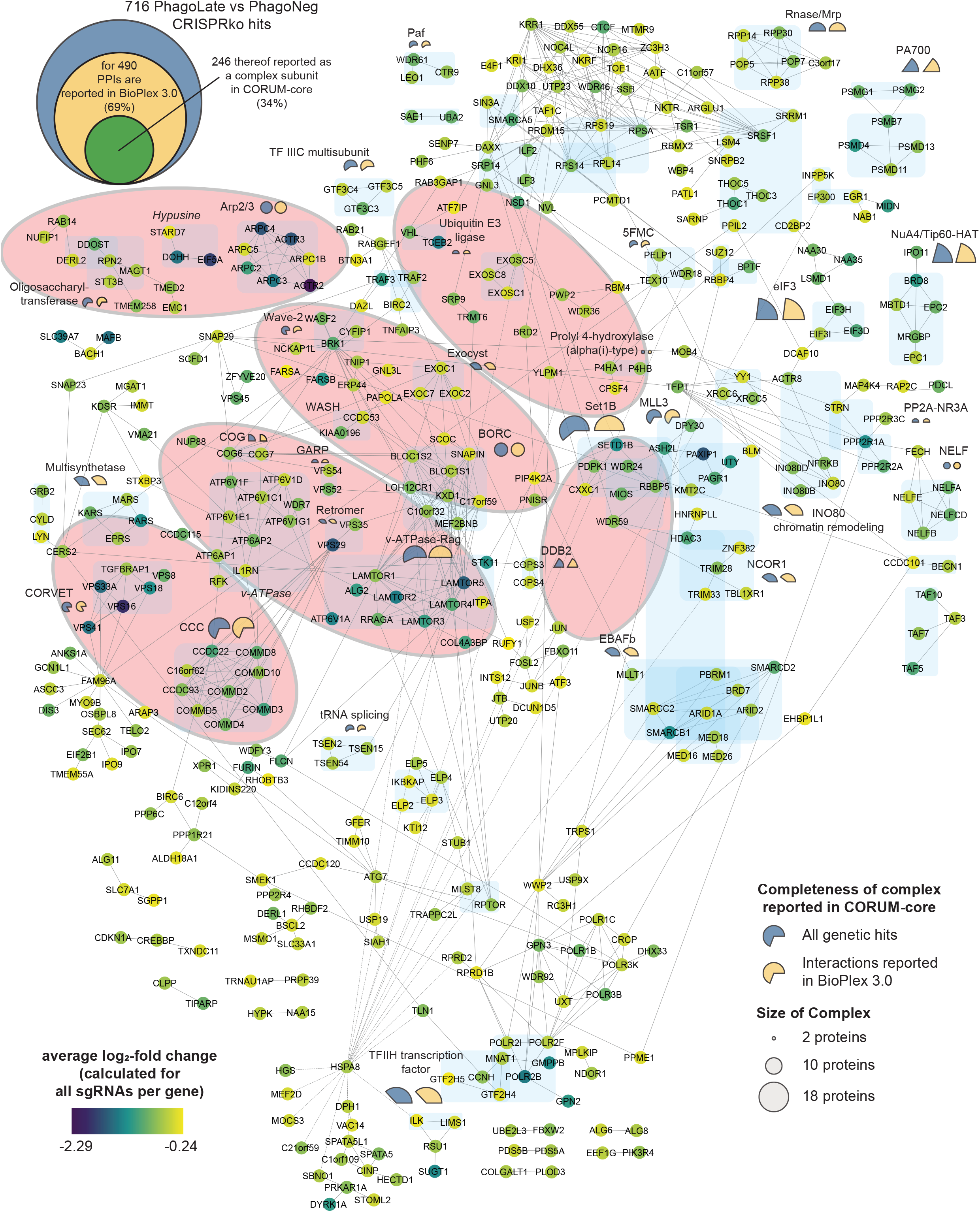
Network of 490 genes that are depleted in the PhagoLate population (d) and have reported protein-protein interactions in BioPlex 3.0. Each circle represents a gene, and the lines represent the reported protein-protein interactions. The color of the circle represents the average log_2_-fold change (aLFC). 246 of the genes are reported as a complex subunit in the CORUM-core database and are highlighted in blue. The percent completeness of these complexes is indicated either with a blue (percent of respective complex covered by all the genetic hits scoring in the screen) or yellow (percent of respective complex covered by all the genetic hits which have interactions reported in the Bioplex 3.0 dataset) pie chart, whereas the size of the circle indicates the total size of the reported complex.

### Mapping the identified phagocytosis genes to the cellular-machinery

The virtually compiled assembly of human cellular protein complexes inferred from collective protein-protein interaction studies is sufficiently advanced to represent a useful framework for modeling the molecular machinery of a cell. If our functionally determined genes were indeed representing important elements in the machinery involved in phagocytosis, they should converge on a limited number of protein complexes. Moreover, these protein complexes should in turn be involved in linked cellular processes and pathways. To test this hypothesis and to probe for physical interactions between the 716 genetic hits, we created a protein-protein interaction network, derived from interactions recorded in the BioPlex 3.0, a high-quality dataset featured by the consistency of scoring across different baits (Huttlin et al., 2021). For 69% of the 716 hits, we could obtain a fairly coherent network with only a few small, isolated subnetworks. For a better interpretation of the network, the color of the circle represents the average log_2_ fold change (aLFC) of each gene (**Figure 2**). To map the proteins linked in the protein network to known molecular machines, we scored membership to the well-characterized human protein complexes of the CORUM-core (Giurgiu et al., 2019), light blue shaded in **Figure 2** (see also **Suppl Table 4-5**). This allowed us to map about a third of the identified phagocytosis modulators onto known biological complexes representing molecular machines (**Figure 2**). Additionally, the completeness of protein-complexes annotated in CORUM-core was analyzed and indicated in percentages with either a blue (percent of complex covered by all the genetic hits scoring in the screen) or yellow pie chart (percent of complex covered by all the genetic hits which have interactions reported in the Bioplex 3.0 dataset), whereas the diameter of the circle indicates the total size of the reported complex (**Suppl Table 6**). Some genes scoring as essential in our screen (**Suppl Table 2**) were expluded as hits and therefore not represented in our network, but may be involved in phagocytosis, such as COG1.

### Actin cytoskeleton nucleation plays an important role in phagocytosis

To ratify our dataset of phagocytic modulators we decided to pick genes for an arrayed validation that displayed the largest aLFC from the identified complexes with the highest completeness (**Figure 3a**). Among these was the Arp2/3 complex (Goley and Welch, 2006). This actin-nucleating machinery has been described to be highly essential for phagocytic cup formation and is also the mechanistic target of our chemical assay control, Cytochalasin D (May et al., 2000). Our genetic screen picked up all subunits of the Arp2/3 complex (ARPC1B, ARPC2, ARPC3, ARPC5, ARPC4, ACTR3, ACTR2), highlighting the high degree of coverage of our experimental set-up. Assessing the reporter uptake of individual *ARPC2* KO cell lines showed significantly decreased phagocytosis (**Figure 3a-b**), validating the findings of the genome-wide pooled screen. Moreover, our screen identified 80% of the members of the Wave-2 complex as hits (WASF2, CYFIP1, ABI1, BRK1, NCKAP1) and most of these had recorded interactors in the BioPlex 3.0 dataset. WASF2 is a scaffolding protein that employs the Arp2/3 complex and has been reported to be essential for the formation of lamellipodia (Oikawa et al., 2004), indicating that our reporter enters macrophages through these structures. Since WASF2 is a central actin-nucleator and scaffolding protein for Arp2/3, without being essential for diverse processes within the cell unrelated to phagocytosis as the Arp2/3 complex, we picked it as the first in-detail validation target (**Figure 3c**). Thus, we derived *WASF2* KO clones from the human monocyte cell line U937 (Liu and Wu, 1992), which is, competent for phagocytosis and amendable to single clone isolation. We used 4 different sgRNAs targeting different exons of *WASF2* (**Figure 3g**). After confirming by immunoblotting the successful knock-out of *WASF2* in these clones (**Figure 3d**), we rescued them either with a *WASF2* cDNA (designed to be resistant against sgRNA 1,2 and 4 but not against sgRNA 3, cutting control that can cut the *WASF2* cDNA, **Figure 3g**) or with a mock cDNA as control. WASF2 is proposed to be a key effector in the life cycle of *Listeria monocytogenes* (Bierne et al., 2005). We took advantage of this to validate our cellular clones. We infected them with Listeria and quantified the total amount of bacteria phagocytosed after 3h hours, as exemplified in **Figure 3e**. Effectively, the *WASF2* knock-out cell lines, that only express the mock control cDNA, harbored lower levels of total phagocytosed bacteria (**Figure 3f**). The phagocytic activity of those cells was then evaluated using the reporter uptake assay. Indeed, cells knocked out for *WASF2*, and only expressing the mock control cDNA, showed a significant ablation for the PhagoLate fraction with a concurrent increase of the PhagoNeg fraction, indicating a decrease in total phagocytosis (**Figure 3h**). At the same time, this defect could be rescued by expressing a *WASF2* cDNA, confirming that the impairment of phagocytosis was caused by the knock-out of *WASF2* (**Figure 3g-h**).

**Figure 3:**
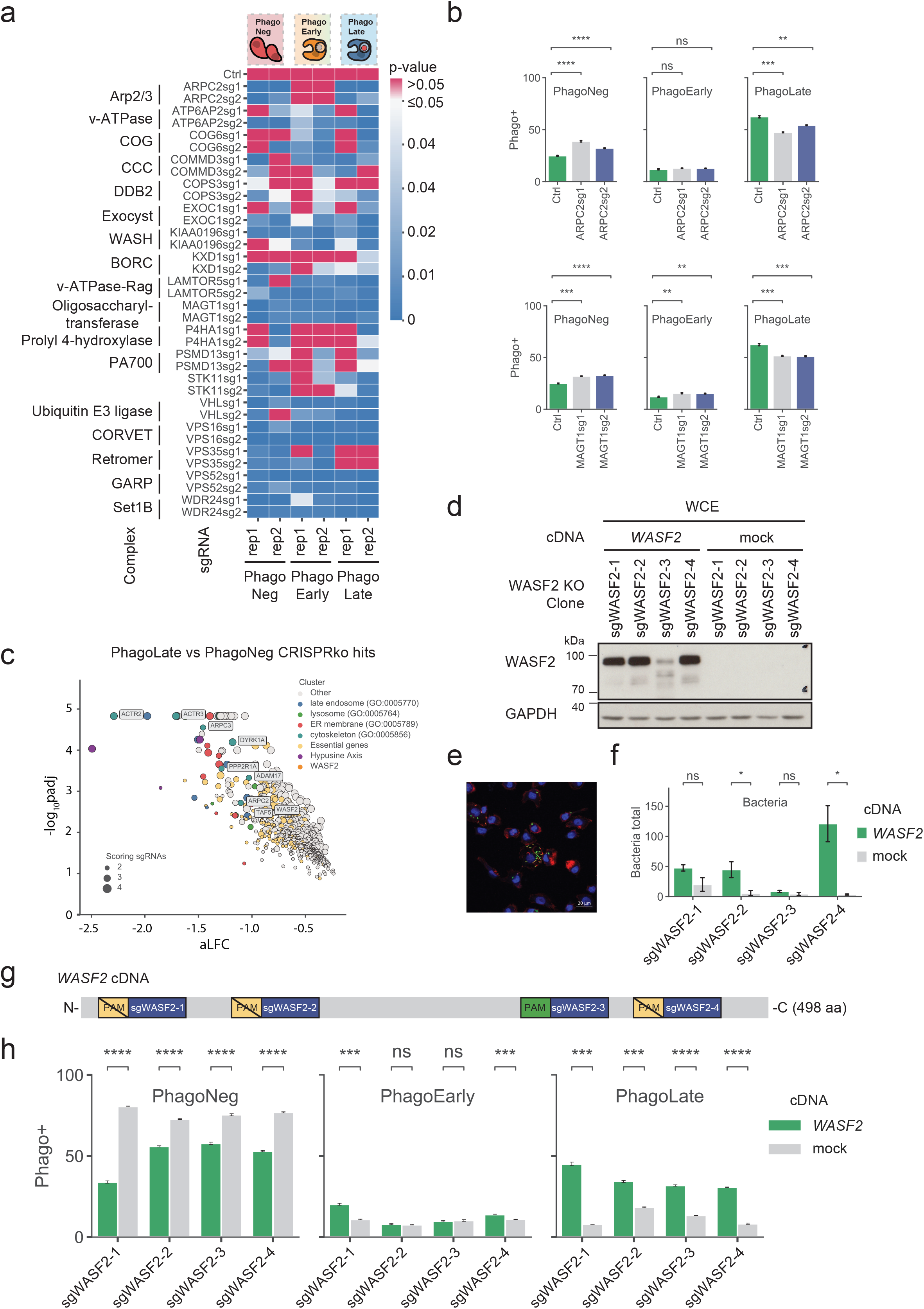
**a**. Heatmap showing the significance of difference of the phagocytosis fractions recorded with phagocytosis assays in the respective knock-out cell lines listed on the y-axis compared to control cell lines, shown in an example in (b). The x-axis shows the quantified population and the replicate number. p-values > 0.05 are considered insignificant and marked in red. Data are mean ± 95% confidence interval from 3 replicates by Welsh’s t test. **b**. Barplot showing the quantification of the PhagoNeg, PhagoEarly, and PhagoLate fraction as examples for validation of the Arp2/3 complex on ARPC2 KO cell lines as well as for the oligosaccharyltransferase complex on MAGT1 KO cell lines. Data are mean ± 95% confidence interval from 3 replicates. **p ≤ 0.01, ***p ≤ 0.001, ****p ≤ 0.0001; by Welsh’s t test. **c**. Volcano plot highlighting WASF2 and genes of the Arp2/c complex scoring within the 716 genes depleted in the PhagoLate population. The x-axis shows the average log_2_-fold change (aLFC) calculated for all sgRNAs per gene against the y-axis, representing the statistical significance as -log_10_padj. The size of the dot indicates the amount of sgRNAs changing significantly for the particular gene. **d**. Western blot showing expression of WASF2 in U937 WASF2 knock-out cell clones, overexpressing either a mock cDNA or a WASF2 cDNA, designed to be resistant to cleavage by sgWASF2-1,2,4 but not sgWASF2-3, as depicted in (f). **e**. Representative image showing the readout used for quantification of the listeria assay. The green color highlights *Listeria monocytogenes*, the red color actin and blue highlights the DAPI staining. **f**. Barplot showing the quantification of the total amount of *Listeria monocytogenes* of cell lines validated in (d). The green color highlights the clonal U937 WASF2 KO cell lines expressing WASF2 and the grey color the same cell lines expressing a mock cDNA as control. Data are mean ± 95% confidence interval from 3 replicates. *p ≤ 0.05, **p ≤ 0.01 by Welsh’s t test. **g**. Schematic representation of the WASF2 cDNA overexpressed in cell lines validated in (d). PAM indicates the protospacer adjacent motif compatible with Cas9 detection of each WASF2 sgRNA, indicated by the blue boxes. **h**. Barplot showing the quantification of the PhagoNeg, PhagoEarly, and PhagoLate fraction of cell lines validated in (d). The green color highlights the clonal U937 WASF2 KO cell lines expressing WASF2 and the grey color the same cell lines expressing a mock cDNA as control. Data are mean ± 95% confidence interval from 3 replicates. ***p ≤ 0.001, ****p ≤ 0.0001; by Welsh’s t test.

Our physical interaction map of genetic hits offers the opportunity to streamline validation by navigating neighboring cellular complexes as a functional framework, instead of choosing randomly among the 716 genes. The underlying assumption is that proximity to a validated node will increase the validation efficiency of the next one, while allowing to derive a functional relationship between the two.

In our network, the Wave-2 complex is connected via BRK1 to the WASH complex (KIAA0196, CCDC53), an endosomal hub responsible for the activation of Arp2/3 (Jia et al., 2010) (**Figure 2**). Knock-out of *KIAA0196* led to a phenotype (significant reduction of reporter uptake) comparable to *WASF2*, confirming the functional connection (**Figure 3a**).

### Lysosomal regulation modulates acidification of the lysosome and affects phagocytosis

Continuing along our “validation path” we observed that the WASH complex is connected multiple times to the BORC complex (BLOC1S1, BLOC1S2, SNAPIN, LOH12CR1, KXD1, C10orf32, C17orf59, MEF2BNB) (Pu et al., 2015), which we identified in our genetic screen with a remarkable 100% completeness, underlining its importance for phagocytosis. Since the BORC complex in our network displays further connections to the vacuolar-ATPase-Rag complex (LAMTOR1, LAMTOR2, LAMTOR3, LAMTOR4, LAMTOR5, ALG2, ATP6V1A, STK11) and is through these further connected to the vacuolar-ATPase protein machinery (ATP6V1F, ATP6V1D, ATP6V1C1, ATP6V1E1, ATP6V1G1, ATP6AP1, ATP6AP2, WDR7, IL1RN) (Kissing et al., 2015), we believe that this subnetwork of hits delineates lysosome regulating functions. ARL8B is another prominent regulator of lysosomal trafficking, that scores in our screen and is part of this subnetwork, but is not represented in our network, as it lacks within BioPlex 3.0 any direct connection with hits identified within our screen (Garg et al., 2011; Pu et al., 2015). We validated this neighborhood by knocking out *LAMTOR5 as* a member of the vacuolar-ATPase-Rag complex and *KXD1*, a member of the BORC complex. Both loss of function cell lines led to a significant reduction in phagocytosis (**Figure 3a**). Interestingly, the target of Bafilomycin A1, vacuolar-ATPase, is also a hit in this subnetwork. In contrast to genetic ablation, acute drug treatment allows revealing the importance of the acidification process of phagocytosis without affecting uptake or cargo shuffling (**Figure 1b**).

### Involvement of the vesicle trafficking machinery in phagocytosis

In our network of functional hits, the BORC complex was also connected to the conserved structural Golgi protein COG6 (Ungar et al., 2002) and through this to four complexes responsible for vesicle trafficking within the cell, such as the exocyst complex (EXOC1, EXOC2, EXOC7) (Katoh et al., 2015), the GARP complex (VPS52, VPS54) (Pérez-Victoria et al., 2010), the retromer complex (VPS29, VPS35) (Haft et al., 2000) and the CORVET complex (VPS16, VPS18, VPS33A, VPS8, TGFBRAP1) (van der Kant et al., 2015). To continue our validation campaign, we generated knock-out cell lines for several subunits of the above-mentioned complexes including EXOC1, VPS52, VPS35, and VPS16. We observed a significant reduction in phagocytosis in all these knock-out cell lines (**Figure 3a**). Interestingly, the knock-out of VPS35, a subunit of the retromer complex, significantly reduced the PhagoNeg and PhagoEarly fraction, but not the PhagoLate fraction, confirming in a functional assessment of the different phagocytosis steps, that VPS35 regulates the uptake of materials to the phagosome and less the later part of phagocytosis. In our network, we could then observe interactions between the retromer complex and all of the subunits of the CCC complex (also known as Commander complex). Within our genetic screen, we covered all 10 in CORUM reported subunits of the CCC complex (COMMD2, COMMD3, COMMD4, COMMD5, COMMD8, COMMD10, CCDC22, CCDC93, VPS29, C16orf62/VPS35L) as single individual hits. The CCC complex is a poorly studied protein assembly that seems to play a role in endosomal cargo retrieval and recycling as well as regulation of nuclear factor-κB (NF-κB) and hypoxia-induced transcription (Cullen and Steinberg, 2018; Laulumaa and Varjosalo, 2021). Through the work presented here, it is possible to ascribe a biological role in phagocytosis to the CCC/Commander complex, likely to be related to the vesicular sorting function of the retromer complex (**Figure 2, Figure 3a**).

### The oligosaccharyltransferase complex is a strong modulator of phagocytosis

Although the oligosaccharyltransferase complex (OST) (RPN2, DDOST, STT3B, MAGT1) forms an independent subnetwork on our map, we decided to further assess its role due to its high completeness in our screen and its novel link to phagocytosis. The OST complex is a core-member of the central machinery for N-linked protein glycosylation. We chose to knock out the magnesium transporter *MAGT1/SLC58A1*, known to be mutated in patients with a disorder disease called X-linked immunodeficiency with magnesium defect, Epstein-Barr virus (EBV) infection, and neoplasia (XMEN) (Ravell et al., 2020). A further hint, that SLC58A1 is involved in phagocytosis, is that *SLC58A1* deficient cells manifested a very strong defect in phagocytosis (**Figure 3a**).

We chose to validate six additional genes that were not part of large molecular complexes but were not previously associated with phagocytosis. Of these hits three showed a less robust effect in the validation process, manifested by variations between individual replicates of phagocytosis assays (*COPS3, P4HA1*, and *PSMD13*). Knock-out of *STK11* mainly negatively affected uptake of material, revealed by decreased PhagoNeg fractions but fewer effects on the PhagoEarly and PhagoLate fractions and Ubiquitin E3 ligase *VHL*, as well as *WDR24*, led to a general defect in phagocytosis (**Figure 3a**).

In summary, of the genes that we assessed individually in phagocytosis assays tracking three different stages, 83% validated effectively (15 out 18). This suggests that, overall, the dataset encompassing 716 genes represents a valuable resource for the molecular characterization of the cellular machinery in the complex phagocytosis process.

### Depletion of the unique posttranslational modification hypusine on eIF5A reduces phagocytosis

Interestingly, the gene with the highest significant depletion in genome-wide phagocytosis screen was Deoxyhypusinesynthase (*DHPS*), scoring together with two other genes (*DOHH* and *eIF5A*) that are all involved in forming the unique post-translational modification hypusine (Park et al., 1981) (**Figure 1d, Figure 4g**). eIF5A is a small ∼17 kDa conserved protein, the mammalian ortholog of the bacterial translation factor EFP (Doerfel et al., 2013), and the only known target of the posttranslational modification hypusine, which is incorporated by an interplay of the two enzymes DHPS and DOHH from the polyamine precursor spermidine (Wolff et al., 2007). Since not only *eIF5A* but also *DHPS* and *DOHH* scored as hits in the screen we concluded that the absence of hypusine must be causative of a phagocytosis defect (**Figure 4a**). To check this, we generated an inducible THP-1 knock-out model for *DHPS* using an inducible CRISPR-Cas9 knockout backbone (TLCV2) (Barger et al., 2019). Furthermore, since there was no commercial antibody available for hypusine we generated a recombinant hypusine-specific antibody based on a sequence published by Zhai and colleagues (Zhai et al., 2016). We validated the antibody by transiently expressing *DHPS, DOHH*, and *eIF5A* (either *eIF5A*-WT or mutated for the hypusine modification site lysine 50) in HEK293T cells. The antibody successfully detected hypusinated eIF5A, while there was no signal detected in cells overexpressing the eIF5A-K50 variant (**Figure 4b**). To further validate the binder we tested it on lysates of THP-1 cells treated with the DHPS inhibitor GC7 (Melis et al., 2017) and observed downregulation of hypusine after 24-72 hours (**Figure 4c**). As GC7 could also act indirectly, we used the inducible *DHPS* knock-out cell models to show that the downregulation of DHPS is sufficient to downregulate hypusine (**Figure 4f**). Having validated our experimental set-up, we measured phagocytosis in both settings. Treating THP-1 cells with GC7 led to a significant drop of the PhagoLate fraction with a parallel rise of the PhagoNeg fraction, suggesting an effect of hypusine on the total rate of phagocytosis (**Figure 4d**). We observed the same effect when we measured phagocytosis after *DHPS* knock-out (**Figure 4h**). Introducing a *DHPS* WT cDNA, designed to be resistant against the sgRNA, could rescue the defect, demonstrating the specific dependency on *DHPS* (**Figure 4e-f, h**). As an additional control of our cell model, we then rescued our *DHPS* knock-out model with *DHPS* cDNAs mutated to strongly reduce spermidine binding (D243A) (Lee et al., 2001). Interestingly, when we rescued our cell model with the D243A mutant *DHPS* cDNA, the cell seemed to compensate for the reduced function of the D243A *DHPS* variant by higher expression levels of D243A compared to other DHPS protein levels (**Figure 4f**). Nevertheless, despite higher expressed levels of the mutant DHPS, we observe less phagocytosis compared to cells rescued with *DHPS* WT cDNA (**Figure 4h**). A clinical study on patients with severe neurodevelopmental disorders reported recessive rare variants in the *DHPS* gene (Ganapathi et al., 2019). Therefore, we wondered if the expression of these mutant DHPS variants would also lead to a defect in our phagocytosis assay. Indeed, when we expressed either the N173S or the ΔY305-I306 patient *DHPS* variant in our *DHPS* knock-out model, we could observe a clear defect in phagocytosis (**Figure 4h**). Expression levels of the mutant DHPS proteins were comparable to the wild type (**Figure 4f**). Thus, *DHPS, DOHH*, and *eIF5A* can be regarded as genes validated for their involvement in phagocytosis.

**Figure 4:**
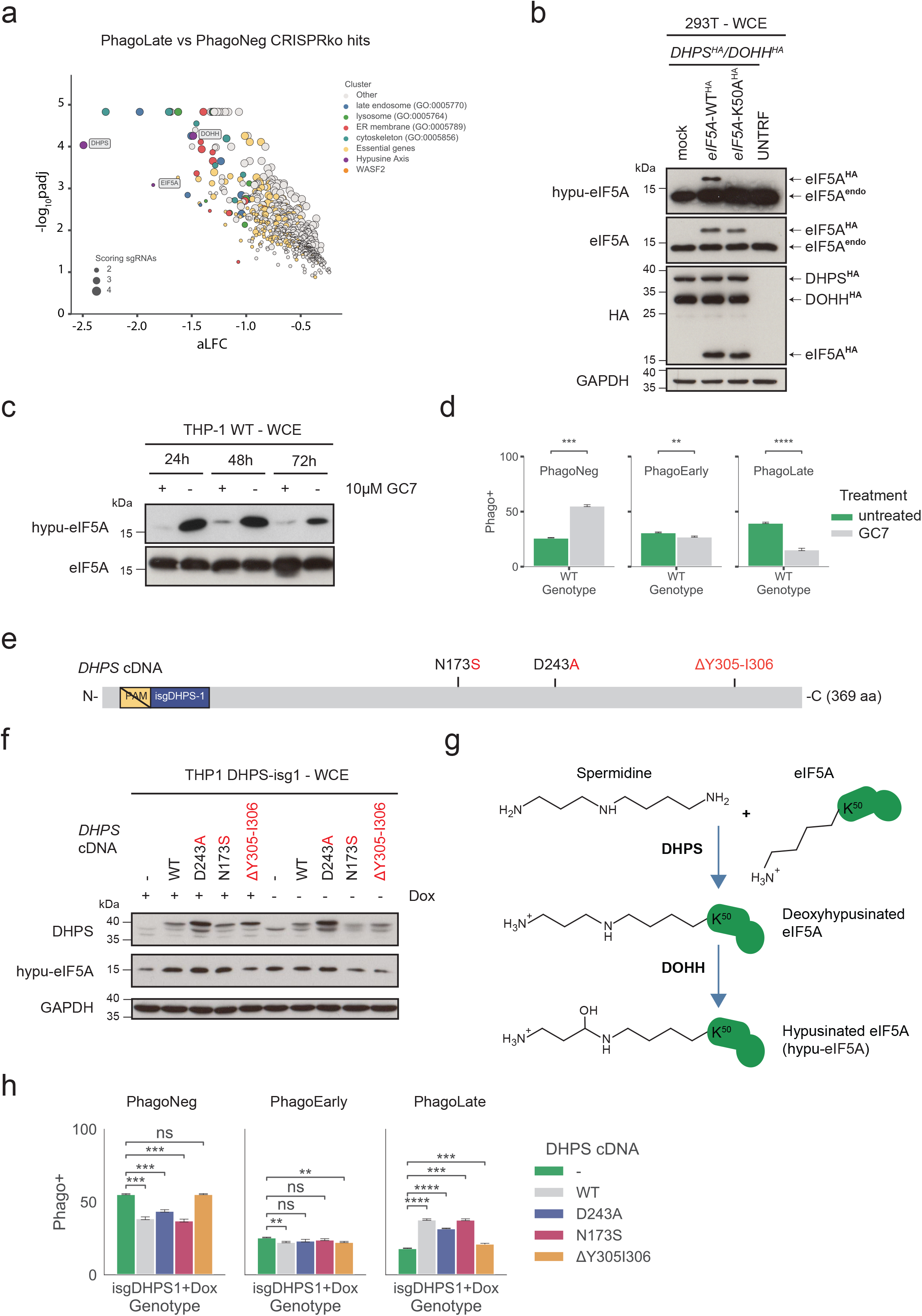
**a**. Volcano plot highlighting the genes of the hypusine axis within the 716 genes depleted in the PhagoLate population. The x-axis shows the average log_2_-fold change (aLFC) calculated for all sgRNAs per gene against the y-axis, representing the statistical significance as -log_10_padj. The size of the dot indicates the amount of sgRNAs changing significantly for the particular gene. **b**. Western blot analysis of hypusine (hypu-eIF5A) in HEK293T cells overexpressing either a mock or C-terminal HA-tagged eIF5A-WT (*eIF5A*-WT^HA^) or eIF5A-K50A mutant cDNA (*eIF5A*-K50A^HA^) as well as a C-termial HA-tagged DHPS cDNA (*DHPS*^HA^) and DOHH cDNA (*DOHH*^HA^) **c**. Western blot analysis of hypusine levels (hpu-eIF5A) in THP-1 WT cells treated with or without 10 µM GC7 at the respective time point (24 – 72 hours). **d**. Barplot showing the quantification of the PhagoNeg, PhagoEarly, and PhagoLate of a phagocytosis assay performed with cells treated with or without 10 µM GC7. Data are mean ± 95% confidence interval from 3 replicates. **p ≤ 0.01, ***p ≤ 0.001, ****p ≤ 0.0001; by Welsh’s t test. **e**. Schematic representation of the cDNA of *DHPS* overexpressed in cell lines validated in (f). PAM indicates the protospacer adjacent motif compatible with Cas9 detection, isgDHPS-1 marks the binding site of the respective sgRNA, and N173S, D243A, and ΔY305-I306 indicate mutations introduced in the corresponding DHPS cDNAs. **f**. Western blot analysis of the expression of WT or mutated DHPS and hypusine levels (hypu-eIF5A) in doxycycline-inducible THP-1 knock-out cells grown either with or without doxycycline. **g**. Schematic showing the key steps and enzymes involved in the hypusine axis. K50 highlights the lysine on eIF5A that undergoes the hypusine modification. **h**. Barplot showing the quantification of the PhagoNeg, PhagoEarly, and PhagoLate of phagocytosis assays performed with the cell lines described in (f). Data are mean ± 95% confidence interval from 3 replicates. **p ≤ 0.01, ***p ≤ 0.001, ****p ≤ 0.0001; by Welsh’s t test.

## Discussion

The FACS-based genome-wide CRISPR/Cas9 knock-out screen, allowed us to identify a large number of genes involved in the regulation and execution of the phagocytosis process. The high number of genes scoring positive in the assay represented a challenge for further validation. We designed a strategy that used an integration with existing protein-protein interaction networks to prioritize which genes to validate. We reasoned that focusing on those molecular machines of the cell, whose components were “genetically hit” multiple times in the screen, would be an expedient to increase the statistical significance of the various cellular functions associated with phagocytosis and thus ensure a higher validation rate. Recently Haney and colleagues undertook a systematic search for phagocytic regulators with a pooled CRISPR screen approach (Haney et al., 2018). They used macrophage differentiated U937 cells and incubated them with magnetic beads conjugated to a series of different substrates such as myelin, zymosan, and sheep red blood cells. In our study, we use the human monocytic cell line THP-1 differentiated into macrophages and a FACS-sorting-based separation approach. We confirmed and expanded their identified regulators as to the importance of the Arp2/3 machinery, Wave-2 complex, and the mTOR-associated Ragulator complex. This was demonstrated by the high genetic coverage of these protein complexes in our dataset. As already shown by Haney et al. for U937 cells, loss of subunits of the actin polymerization machinery is detrimental to phagocytosis in THP-1 cells. Several datasets collected in screening efforts for host regulators in the context of *Salmonella* (Yeung et al., 2019), *Legionella pneumophila* (Jeng et al., 2019) and *Staphylococcus aureus* (Lindner et al., 2021) pathogenesis confirmed the importance of the actin polymerization machinery. This indicates that our pathogen-agnostic reporter approach used in this screen successfully mapped the cellular routes taken by pathogens. Furthermore, we identified several members of the exocyst complex to be required for phagocytosis. Previous studies on the interaction of Cdc42 with the exocyst complex, found to be essential for promoting phagocytosis (Mohammadi and Isberg, 2013), helped us to interpret the scoring of the exocyst complex in our screen. As Cdc42 is a Rho GTPase central for actin dynamic (Bonfim-Melo et al., 2018), we can now position the exocyst complex close to the family of actin regulators in our hit reconstructed protein interaction network (**Figure 2**). Despite affecting phagocytosis in our screen, *Cdc42* has very high general essentiality and therefore was scored correctly as an essential gene rather than a hit in our screen (**Suppl Table 2**). This case demonstrated the well-known drawback of knock-out screens for specific processes and subsequent readouts, which typically fail to identify highly essential genes. In the future, the issue could be fixed by using a loss of function system that reduces and does not abolishe gene functions (such as CRISPR interference screens) (Gilbert et al., 2014; Rossi et al., 2015; Rousset et al., 2018).

Our study also confirmed the importance of the positive regulation of mTOR signaling for phagocytosis (Haney et al., 2018). A finding that was also revealed by the identification of RagA (*RRAGA*), as a regulator of microglial phagocytic flux in a zebrafish screen (Shen et al., 2016). Remarkably, our screen identified all five members of the Ragulator complex (LAMTOR1, LAMTOR2, LAMTOR3, LAMTOR4, and LAMTOR5) as well as *RRAGA*. In addition, we found the full BORC complex, a machinery responsible for positioning lysosomes. The BORC complex is reported to interact with Ragulator, a complex that negatively regulates the activity of BORC in response to amino acid starvation sensed by SLC38A9, a critical regulator of lysosomal function that we had previously identified (Filipek et al., 2017; Rebsamen et al., 2015; Wyant et al., 2017). The role of BORC in phagocytosis can potentially be explained by a recent preprint showing the importance of TORC1, BORC, ARL-8, and kinesin-1 for vesiculation of the phagolysosome and especially the role of BORC as a promoter for phagolysosomal degradation (Fazeli et al., 2022).

Our screen also highlighted the importance of phagocytosis of coordinated cargo transport through the cell. This requires the trafficking and fusion of endosomes, controlled by Rab-GTPases, SNARE proteins, and multi-subunit tethering complexes (Bröcker et al., 2010). Using our dataset to identify the parts of this canonical vesicle trafficking machinery of most relevance for phagocytosis, we observed that large parts of the CORVET and HOPS complexes, machinery that coordinate together with Rab5 and Rab7 (*RAB5C* and *RAB7A* are hits as well) the fusion of the endosome with the lysosome, score as strong phagocytic regulators. Notably, VPS16, the shared subunit of both the CORVET and HOPS complex, showed the strongest depletion in our screen among the above-mentioned regulators (van der Kant et al., 2015). Likewise, we identified large parts of the GARP complex, which together with the COG complex forms the Golgi machinery associated with phagocytosis (Pérez-Victoria et al., 2010), and large parts of the retromer complex, an apparatus responsible for the recycling of organelles in the cell (Haft et al., 2000).

*MAGT1/SLC58A1*, a gene whose loss-of-function mutations cause X-linked immunodeficiency with magnesium defect (XMEN; OMIM: 300853) (Watson et al., 2022), significantly scored together with other parts of the OST complex as phagocytosis regulators in our screen and was previously found as host cell regulators of *Legionella pneumophilia* (Jeng et al., 2019). Either MAGT1/SLC58A1 or its paralog TUSC3/SLC58A2 can assist the STT3B-OST complex during post-translational N-glycosylation in the endoplasmic reticulum (Harada et al., 2019; Ravell et al., 2020). However, since *TUSC3* is not expressed in immune cells, THP-1 cells rely only on *MAGT1* (Watson et al., 2022) and indeed our screen scored only *MAGT1* but not *TUSC3* as a phagocytic regulator. As noted MAGT1/SLC58A1 is also a transporter of magnesium ions (Mg2+), a metal that is recognized recently as being crucial for the regulation of immune response (Maier et al., 2021). Therefore, both functions of SLC58A1, its facilitating role in the STT3B-OST complex, as well as the magnesium transport function, could play a crucial role in phagocytosis. Since we additionally identified the accessory proteins (DDOST and RPN2) and the catalytic subunits (STT3A and STT3B) of the OST complex as hits, we conclude that N-linked glycosylation plays an important general role in phagocytosis and consequently speculate that dysregulated phagocytosis might be a factor in XMEN disease (Ravell et al., 2020). How exactly N-linked glycosylation affects phagocytosis will require dedicated investigations.

Beyond the hits described so far, numerous uncharacterized genes, that had never been previously linked to phagocytosis, scored in our screen. This valuable resource is included in the dataset associated with this study (**Suppl Table 1**). A detailed validation of identified genes will undoubtedly represent a worthwhile follow-up of the work presented here.

To our surprise, however, the top depleted gene in our screen was *DHPS*, along with its two interacting proteins DOHH and eIF5A, which both scored with very high significance. eIF5A is one of the 20 most abundant proteins in proliferating cells (Hukelmann et al., 2016) and therefore it is not unexpected that it has been implicated in many cellular functions such as translational elongation (Saini et al., 2009), nucleoplasmic transport (Rosorius et al., 1999), cell proliferation, and as a modulator for mRNA decay (Valentini et al., 2002). DHPS generates the post-translational modification deoxyhypusine by the conjugation of the aminobutyl moiety of spermidine on the lysine of eIF5A which is then matured to hypusine by DOHH (Park et al., 1981). Over the years, elegant work led to the structure of the nuclear shuttle for hypusinated eIF5A (Xpo4) (Aksu et al., 2016), the description of RNA binding properties for hypusinated eIF5A (Xu et al., 2004), and reports on patients with mutations in *eIF5A* (Faundes et al., 2021), *DHPS* (Ganapathi et al., 2019) or *DOHH* (Ziegler et al., 2022) that all showed neurodevelopmental phenotypes. Although until recently (Lindner et al., 2021) the hypusine axis was never identified in the context of phagocytosis, several previous studies suggested its implication in vesicular trafficking, cytoskeleton organization, and immune cell regulation, all processes highly relevant for phagocytosis. Already 1952, spermine was found as an anti-mycobacterial natural agent (Hirsch and Dubos, 1952) and later as an inhibitor for cytokine synthesis in human mononuclear cells (Zhang et al., 1997). Yeast studies found a synthetic lethal interaction between *eIF5A* and *YPT1* (the yeast ortholog of mammalian rab genes, coordinators of endocytosis) (Clague, 1998; Frigieri et al., 2008) and the regulators for yeast cell polarity and actin nucleation (*PKC1, ZDS1, GIC1* (human ortholog *CDC42*) as well as *PCL1* and *BNI1*) as suppressors for *eIF5A* depletion (Zanelli and Valentini, 2005). Like the bacterial ortholog EFP (Doerfel et al., 2013), eIF5A was found to assist the translation of polyproline stretches in mammalian proteins (Barba-Aliaga et al., 2021; Gutierrez et al., 2013; Ude et al., 2013) that are especially enriched in proteins of the actin/cytoskeleton, RNA splicing/turnover, DNA binding/transcription, cell signaling (Mandal et al., 2014) as well as in collagen. And while the DOHH homolog nero and eIF5A, showed to be required for autophagy in drosophila (Patel et al., 2009), eIF5A was found in human cells to regulate the translation of TFEB and therefore also regulate autophagy (Zhang et al., 2019). In addition, eIF5A was shown to assist with the co-translational translocation of proteins in the ER (Rossi et al., 2014), and ablation of eIF5A triggers ER stress in mammalian cells (Mandal et al., 2016). Recently, the hypusine axis was described in the context of macrophage respiration (Puleston et al., 2019), and as the effector of macrophage inflammatory state in adipose tissue (Anderson-Baucum et al., 2021). Finally, mice lacking hypusine are more susceptible to *Heliobacter pylori* and *Citrobacter rodentium* infections (Gobert et al., 2020).

Since we identified the whole hypusine axis and eIF5A as the unique carrier of hypusine, we hypothesized that the hypusinated form of eIF5A must play a significant role in phagocytosis. Our experiments confirmed that the absence of hypusine, either by blocking DHPS, the key enzyme for hypusination, pharmacologically with GC7 or by a knock-down of DHPS with Cas9 indeed led to reduced phagocytosis. Remarkably, we could further show that this defect in phagocytosis also occurred when we replaced the endogenous version of DHPS in THP-1 cells with recessive rare variants of mutated DHPS found in patients (N173S and ΔY305-I306) with neurodevelopmental disorders or with a DHPS variant mutated in the predicted binding site of spermidine (D243A).

In summary, our screen functionally outlined the basic phagocytic machinery to involve several hundred gene products organized in at least two dozen protein complexes. Phagocytosis as a long and complex process and functionally requires the contribution of several subprocesses, such as cytoskeletal mechanical and membrane uptake machinery involved in engulfing and internalizing the foreign material in the phagosome, trafficking of the phagosome, fusion with the lysosome followed by acidification and clearance and recycling to the plasma membrane. Here, publicly available protein-protein interaction data and complex annotation allowed us to assign genes to cellular functions, at least in those instances where the annotation of cellular protein complexes was unequivocal. Our work offers an improved understanding of how cargo travels through the endo lysosomal system and we anticipate that this could in turn direct therapeutics better to their proposed site within the cell, such as antibody-drug conjugates (Tsui et al., 2019) or Cas9 (Lino et al., 2018). In addition, it might help to identify new targets for interventions in diseases caused by dysfunctional autophagy or endo-lysosomal systems such as the neurodegenerative diseases Alzheimer’s disease, Parkinson’s disease, or Huntington’s disease. Ultimately, we believe that mining and integrating publicly available protein-protein interaction data with reporter-based genome-wide genetic screens showcased in this study offers a powerful and widely applicable approach for genetically mapping complex biological processes.

## Materials and Methods

### Cell lines

THP-1, U937, and HEK293T cells were purchased from ATCC. THP-1 and U937 were maintained in RPMI1640. HEK293T were maintained in Dulbecco’s modified Eagle’s medium (DMEM). All media were supplemented with 10% fetal bovine serum (FBS) and antibiotics (100 U/mL penicillin, 100 mg/mL streptomycin), all either from Gibco or Sigma. Cell lines were grown at 37°C in 5% CO_2_.

### Phagocytosis assay

Phagocytosis assay was conducted as previously described (Colas et al., 2014). Reporter beads were generated by opsonization of dual-colored 1.75 µm latex beads (Fluoresbrite carboxylated 1.75 µm microspheres (yellow-green: 441-nm excitation, 486-nm emission, Polyscience, cat no. 17687-5)) in 50% human male AB serum in PBS for 16 h at 4°C under permanent rotation. Then beads were washed twice with PBS and labeled with 2 µg/mL pHrodo-Red SE in PBS (Thermo Scientific, cat no. P36600), for 30 min at room temperature under constant agitation. Reporter beads were then washed with PBS and adjusted to a final concentration of 1 × 10^9^ beads per mL in PBS.

For phagocytosis assays that were performed on individual cell lines, PMA-differentiated cells (THP-1 or U937) were seeded on 12-well cell-culture coated dishes (1 × 10^6^ cells per well). Reporter beads were then added to the cells at an MOI of 10 for 3 hours. Cells were then transferred onto ice, washed three times with ice-cold PBS, detached by scraping with a soft rubber cell scraper and analyzed on an LSR Fortessa II cytometer controlled with FACSDiva (BD), analyzed with FlowJo software (v.10), and visualized with Python project version 3.8.8 (Python Software Foundation. Available at http://www.python.org) with pandas (Reback et al., 2022) (1.2.4), numpy (1.20.1), matplotlib (Hunter, 2007) (3.3.4), seaborn (Waskom, 2021) (0.11.1) and statannotations (0.4.4).

For the Cytochalasin D (Enzo, cat no. BML-T109-0001) and Bafilomycin A1 (Enzo, cat no. BML-CM110-0100) control assays, cell media was supplemented with the compound 30 min before the addition of the reporter beads and added during the whole assay at final concentrations of 8 µM and 200 nM respectively.

For the GC7 (EMD Millipore, cat no. 259545) treated phagocytosis assays, cell media was supplemented during the entire differentiation and seeding period with 10 µM of the compound.

### Genome-wide CRISPR-Cas9/KO screening and cell sorting

For screening, the genome-wide CRISPR-Cas9/KO Toronto Knockout version 3 library from Hart and team (Hart et al., 2017) (Addgene no. 90294), cloned into the 1 vector system (lentiCRISPRv2 carrying Cas9 and sgRNA expression on the same vector) was used. The library consists of 70948 guides, targeting 18053 protein-coding genes with 4 sgRNAs each as well as 142 control sgRNAs. Viral particles were produced by transient transfection of HEK293T cells with the library and packaging plasmids pMD2.G (Addgene no. 12259) as well as psPAX2 (Addgene no. 12260) using PEI (Sigma). 8 hours post-transfection the medium was exchanged to RPMI1640 supplemented with 10% FBS. 72 hours after transfection the viral supernatant was collected, filtered (0.45 µm), and stored at -80 °C until further use. THP-1 cells were infected in quadruplicates with this viral supernatant (supplemented with 8 µg/mL protamine sulfate) at a library coverage of 3000x and a multiplicity of infection of around 0.3, allowing a single viral integration event per cell. Infected cells were then selected with 2 µg/mL puromycin for 4 days, followed by two days of growth in puromycin-free medium, and afterwards differentiated with 10 nM PMA (Sigma) to induce a macrophage-like phenotype. Phagocytosis assay was performed as described in the section “Phagocytosis assay” using dual-colored opsonized latex beads. 80 million cells (∼1000x coverage) were used as input for each phagocytosis assay. All cells were then sorted on a BD FACS Aria II, collecting Phagocytosis positive (PhagoLate) and phagocytosis negative (PhagoNeg) populations. Genomic DNA was extracted using the QIAamp DNA Mini Kit (Qiagen) and the sgRNA-containing cassettes were amplified with a one-round PCR approach following the method of Joung and team (Joung et al., 2017). Amplified samples were then sequenced on a HiSeq2000 (Illumina) at the Biomedical Sequencing Facility (BSF at CeMM, https://biomedical-sequencing.at) and analyzed by a custom analysis pipeline (see below).

### Analysis of CRISPR screens

Sequences of sgRNAs were extracted from NGS reads, matched versus the original sgRNA library index, counted using an in-house Python script and their abundance assessed using a two-step differential approach. First differential abundance of individual sgRNAs was calculated using DESeq2 (Love et al., 2014). Afterward, sgRNAs were sorted by adjusted p-value and aggregated using the gene set enrichment algorithm (Korotkevich et al. 2016; Kuleshov et al. 2016; Subramanian et al. 2005).

### Protein-Protein interaction mapping of genetic hits

To probe for physical interactions between the 716 top hits of the genetic screen, a protein-protein interaction (PPI) network was generated. The protein interactions were extracted from BioPlex 3.0 (for HEK293T and HCT116, downloaded on 20.04.2022) (Huttlin et al., 2021), whereas only the protein interactions between the genes identified as high scoring within the genetic screen were considered. For the extraction of protein interactions all bait-prey interactions reported in BioPlex 3.0 were considered, resulting in a PPI network with 490 proteins (69 % of all genetic hits) with 726 interactions. The network was further organized into protein complexes and modules by annotating all proteins within the network with complexes reported in CORUM-core (the comprehensive resource of mammalian protein complexes, Corum 3.0 current release, downloaded on 20.04.2022) (Giurgiu et al., 2019). Within CORUM homo dimers were removed. 246 proteins (50% of all) in the interaction network were reported as a subunit in protein complexes. Network visualization and ordering were conducted in Cytoscape (v.3.8.0). As a first step all duplicated edges were removed as the directionality obtained by reciprocal AP-MS is not of relevance for the visualization of protein complexes within the network. The interactions were grouped into functional modules by following the association to protein complexes, whereas if a protein mapped to multiple complexes, which is often the case as the CORUM-core contains complex identifiers with partially overlapping subunits, the complex with the highest completeness was considered. The selection of interactions and mapping of complexes were prepared with the statistical software R (v.4.1.3).

### Plasmids and cloning

Knock-out cell lines were generated using the lentiCRISPRv2 CRISPR-Cas9 knock-out vector (Addgene no. 52961). Inducible knock-out cell lines were produced with the TLCV2 CRISPR-Cas9 backbone (Addgene no. 87360) that includes a doxycycline-inducible Cas9-2A-eGFP cassette. In short, for each gene sgRNAs were designed using the CHOPCHOP prediction tool (Labun et al., 2019). Oligos, containing BsmBI compatible overhangs were annealed and cloned into lentiCRISPRv2 using Golden Gate assembly.

Cloned oligonucleotides were as follows (5’ to 3’ orientation):

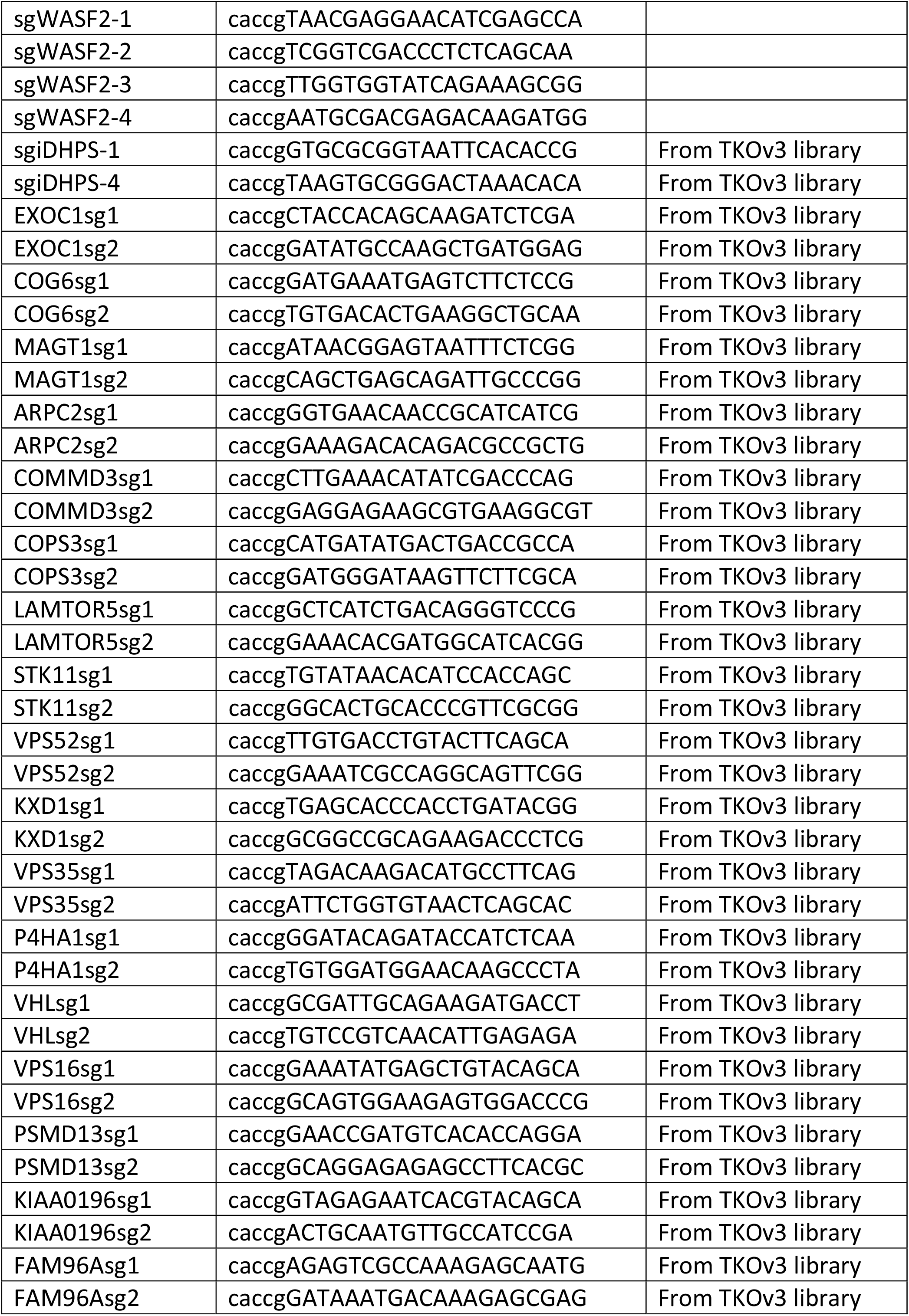
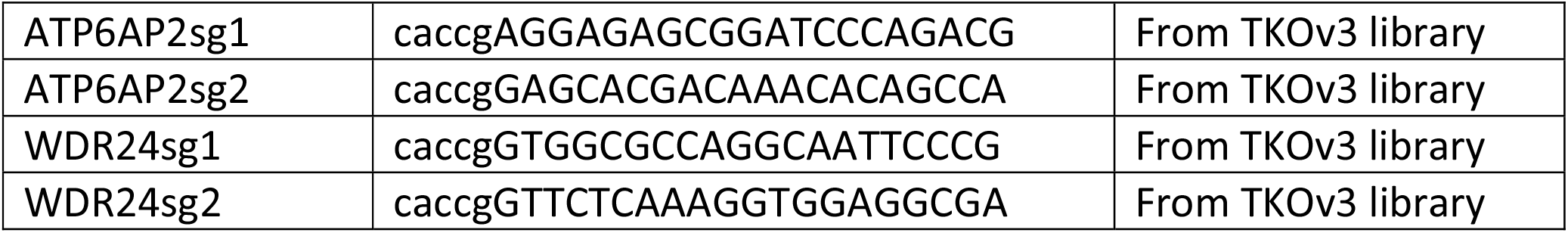

Non-codon optimized cDNA for WASF2 was synthesized and cloned into the ENTR-Twist-Kozak vector by Twist. Mutations in the PAM site of WASF2-sgRNA1, WASF2-sgRNA2, and WASF2-sgRNA4 were introduced to make cDNA resistant against those sgRNAs, using NEB Q5 site-directed mutagenesis kit. After sequence verification, the cDNA was cloned into pRRL-based lentiviral expression plasmids generated in our lab (Bigenzahn et al., 2018), which contain a STOP codon and a BlastR resistance cassette, using the Gateway cloning system (Thermo Fisher Scientific).

DHPS (HsCD00295718, Harvard Medical School plasmid repository), eIF5A (HsCD00041224, Harvard Medical School plasmid repository), and DOHH (HsCD00372432, Harvard Medical School plasmid repository) plasmids were received, and point mutations to make the cDNA resistant against sgRNAs or patient variant mutations were introduced, using the NEB Q5 site-directed mutagenesis kit. They were cloned into the same pRRL-based lentiviral expression plasmids as mentioned above, which contained this time however either an HA-tag or STOP codon on the C-terminus and again a BlastR resistance cassette, using the Gateway cloning system (Thermo Fisher Scientific).

Rescue experiments were conducted by using the cDNAs resistant to the sgRNAs targeting the endogenous genes.

### Lentiviral transduction

Knock-out cell lines and cDNA overexpression cell lines were generated by lentiviral transduction. Therefore, HEK293T were transfected with the respective lentiviral vectors and packaging plasmids pMD2.G (Addgene no. 12259) as well as psPAX2 (Addgene no. 12260) using PEI (Sigma). 16 hours post-transfection the medium was exchanged to RPMI1640 supplemented with 10% FBS. 72 hours after transfection the viral supernatant was collected, filtered through 0.45 µm polyethersulfone filters (GE Healthcare), supplemented with 8 µg/mL protamine sulfate, and directly added to the respective cells. 24 hours after infection, the medium was exchanged, and 48 hours after infection cells were selected with the respective antibiotics.

### Listeria infection assay

*Listeria monocytogenes* strain 10403S (PMID: 3114382) was grown in Brain-Heart infusion medium overnight at 37°C. The infection assay was performed on individual PMA-differentiated U937 cell clones, knocked out for WASF2 and rescued either with a WASF2 cDNA (depicted in **Figure 3f**) or with a mock cDNA as control. A day before infection, 3.0×10^5^ cells were seeded on sterile glass coverslips laid at the bottom of the wells of a 24-well plate in RPMI medium supplemented with 10% FCS and containing no antibiotics. On the day of the infection, bacteria were washed with PBS and incubated at a concentration of 8.0×10^9^ b/mL with 50 µM of the green fluorophore carboxyfluorescein succinimidyl ester (CFSE) for 20 min at 37°C under shaking. Bacteria were then washed with PBS and used to infect the U937 cells at a multiplicity of infection (MOI) of 10. After one hour of infection, the cells were gently washed twice with warm cell culture medium. For the next two remaining hours, the cells were incubated in cell culture medium containing 20 µg/mL of gentamycin to kill the remaining extracellular bacteria. The infection was terminated by washing the cells with PBS and fixing them with 4% paraformaldehyde (Alfa Aesar) for 15 min. Cells were then washed with PBS and permeabilized with 0.1% of Triton X-100 (Roth) in PBS for 5 min. After another washing step with PBS, cells were blocked with 2% FCS in PBS for 30 min. Actin was then stained by incubating the cells with Phalloidin coupled to the fluorophore Alexa647 (Life Technology). Finally, cells were washed with PBS and mounted in Prolong Diamond containing DAPI (Molecular Probe) to stain genomic DNA. All staining steps were carried out at room temperature in the dark. Infected cells were imaged with the confocal microscope LSM 700 (ZEISS) using 405 nm, 488 nm or 639 nm laser lines. Images were processed by the ZEN Software 2009 (ZEISS). Thanks to the gentamycin protection assay, all bacteria detected were considered as phagocytosed. Green bacteria were considered still enclosed inside the phagolysosome whereas yellow bacteria (CFSE and actin colocalized) had successfully escaped the phagolysosome. Quantified bacteria Bacteria was analyzed and visualized with Python project version 3.8.8 (Python Software Foundation. Available at http://www.python.org) with pandas (Reback et al., 2022) (1.2.4), numpy (1.20.1), matplotlib (Hunter, 2007) (3.3.4), seaborn (Waskom, 2021) (0.11.1) and statannotations (0.4.4).

### Gene set enrichment analysis

Enrichment analysis for “Cellular components” GO terms and biological processes GO terms was performed via GSEA (Subramanian et al., 2005) and Enrichr (Kuleshov et al., 2016) using the Python package GSEApy (version 0.10.8, https://github.com/zqfang/GSEApy). Gene sets are described in the corresponding figure legends.

### Cell lysis and immunoblotting

Cell pellets were lyzed in RIPA lysis buffer (20 mM TRIS/HCl pH 7.5, 150 mM NaCl, 1 mM EDTA, 1 mM EGTA, 1% NP40, 1% Sodium deoxycholate, 10 mM NaF, 0.1% SDS, supplemented with EDTA-free protease inhibitor (Roche) for 15 min on ice, followed by 20600g at 4°C for 15 min to separate from cellular debris. Protein concentration was then measured with BCA (Pierce) or Bradford (BioRad), Laemmli buffer was added, and samples were separated by SDS-PAGE and transferred to nitrocellulose membranes (GE Healthcare). Then membranes were blocked for unspecific binding in 5% milk in TBS-T followed by incubation with indicated antibodies. After incubating in peroxidase coupled secondary antibodies, the membranes were developed using the ECL western blot system (Thermo Scientific).

### Antibodies list

The primary antibodies used for western blot were: Hypusine (generation see below, 1:20000), eIF5A (611976, BD Biosciences, 1:20000), DHPS/DHS (ab190266, abcam, 1:1000), DOHH (376929, B-12, SantaCruz, 1:500), HA (3724, C29F4, 1:500), for WASF2 the WAVE-2 (3659, D2C8, CellSignaling, 1:1000), GAPDH (47724, 0411, SantaCruz, 1:2000).

The secondary antibodies used were: anti-rabbit HRP (111-035-144, Jackson ImmunoResearch, dilution 1:10000), anti-mouse HRP (115-035-003, Jackon ImmunoResearch, dilution 1:10000).

A custom rabbit anti-Hypusine antibody was designed based on a 2016 published sequence of context-independent anti-hypusine antibodies (Zhai et al., 2016) (**Suppl Fig 3a**), synthesized, expressed, and purified with Genscript (**Suppl Fig 3b**). In short, the synthesized sequence was cloned into pcDNA3.4 and plasmid DNA prepared. Expi293F cells, grown in serum-free Expi293F Expression Medium (Thermo Fisher Scientific), at 37°C, 8% CO_2_ on an orbital shaker, were then transfected according to manuals. On day 6 post-transfection cell culture supernatant was collected and used for purification. Therefore, the supernatant was centrifuged and filtered, and then loaded and purified on Monofinity A Resin Prepacked Columns. Eluted antibody was then puffer exchanged, sterilized via a 0.22 µm filter, and stored in PBS pH 7.2 at -80°C. The concentration was determined with A280 as 4.74 mg/mL. Specificity was validated by western blot in HEK293T cells overexpressing DHPS/DOHH and either an eIF5A-WT sequence or an eIF5A-K50 variant (**Figure 4b**), as well as in THP-1 cells treated with 10 µM of the DHPS inhibitor GC7 (**Figure 4c**).

### Gene expression analysis

Libraries compatible with Illumina sequencing were prepared using the QuantSeq 3’ mRNA-seq Library prep kit (Lexogen) according to the manufacturer’s instructions. Samples were multiplexed and then sequenced on a HiSeq 4000 (Illumina) at the BSF at CeMM. Raw sequencing reads were demultiplexed and barcode, adaptor, and quality trimming were performed with cutadapt (https://cutadapt.readthedocs.io/en/stable/). Quality control was then performed using FastQC (http://www.bioinformatics.babraham.ac.uk/projects/fastqc/). The remaining reads were mapped to the GRCh38/ h38 human genome assembly using genomic short-read RNA-seq aligner STAR v.2.5. More than 98% of mapped reads in each sample could be obtained with 70–80% of reads mapping to a unique genomic location. End Sequence Analysis Toolkit was used to quantify the transcripts. We carried out differential expression analysis using independent triplicates with DESeq2 (v.1.24.0) on the basis of read counts. Exploratory data analysis and visualizations were performed in R-project v.3.4.2 (Foundation for Statistical Computing, https://www.R-project.org/) with Rstudio IDE v.1.2.1578, ggplot2 (v.3.3.0), dplyr (v.0.8.5), readr (v.1.3.1) and gplots (v.3.0.1).

## Data Availability

Genomics datasets are provided in **Suppl Tables 1-2**. The transcriptomics dataset is attached in **Suppl Table 3**. Protein-protein interaction data mined from BioPlex 3.0 and CORUM-Core specifically for this study are provided in **Suppl Tables 4-6**.

## Acknowledgements

We thank all members of the Superti-Furga laboratory for feedback, discussions and reagents, the Max Perutz Labs FACS Facility for sorting, the Biomedical Sequencing Facility (CeMM/Medical University of Vienna) for next-generation sequencing. In particular, we thank Andrea Garofoli for advice on statistics. We thank Kai-Chun Li and Andreas Angermayr for scientific discussions and critical reading. CeMM and the Superti-Furga laboratory are supported by the Austrian Academy of Sciences. We further acknowledge receipt of third-party funds from the European Research Council (ERC) Advanced Grant 695214 GameofGates (P.E., V.S., G.S.-F.).

## Author Contributions

P.E. and G.S.-F. conceived the study. P.E. and V.S performed and analyzed the genetic screen. P.E., F.F., D.S., L.H., A.S. and B.N. performed experiments. P.E. and G.S.-F. wrote the manuscript with contributions from all authors.

## Supplementary Figures legends

**Figure S1:**
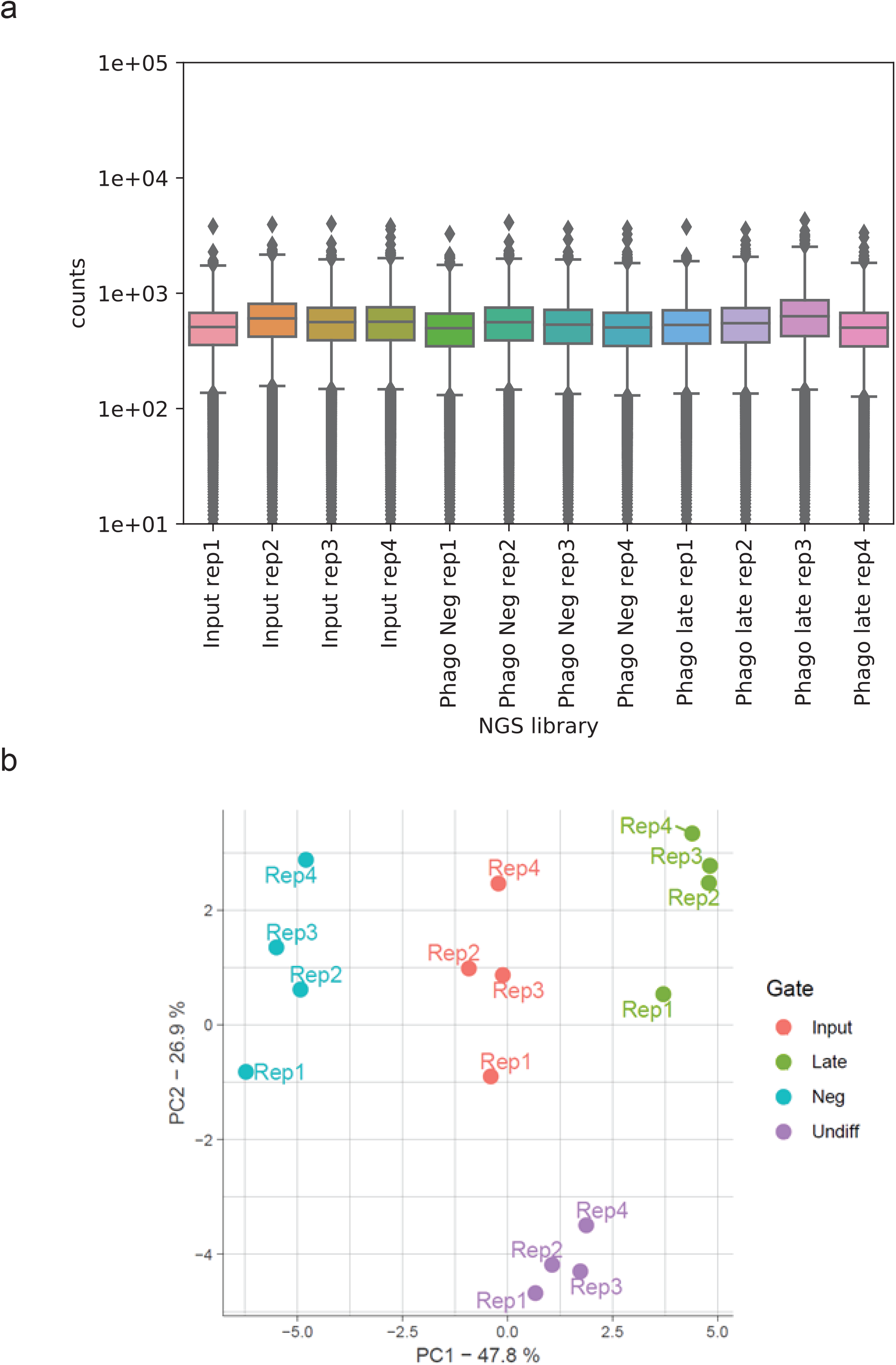
**a**. Read counts for the next generation sequencing (NGS) samples used in this study. Box plots showing the median and the 25/75 percentile. **b**. PCA of counted sgRNAs from NGS reads in the NGS libraries generated from the amplified genomic DNA isolated from the PhagoNeg and PhagoLate populations as well as from the cellular input material and undifferentiated cells, conducted in quadruplicates.

**Figure S2:**
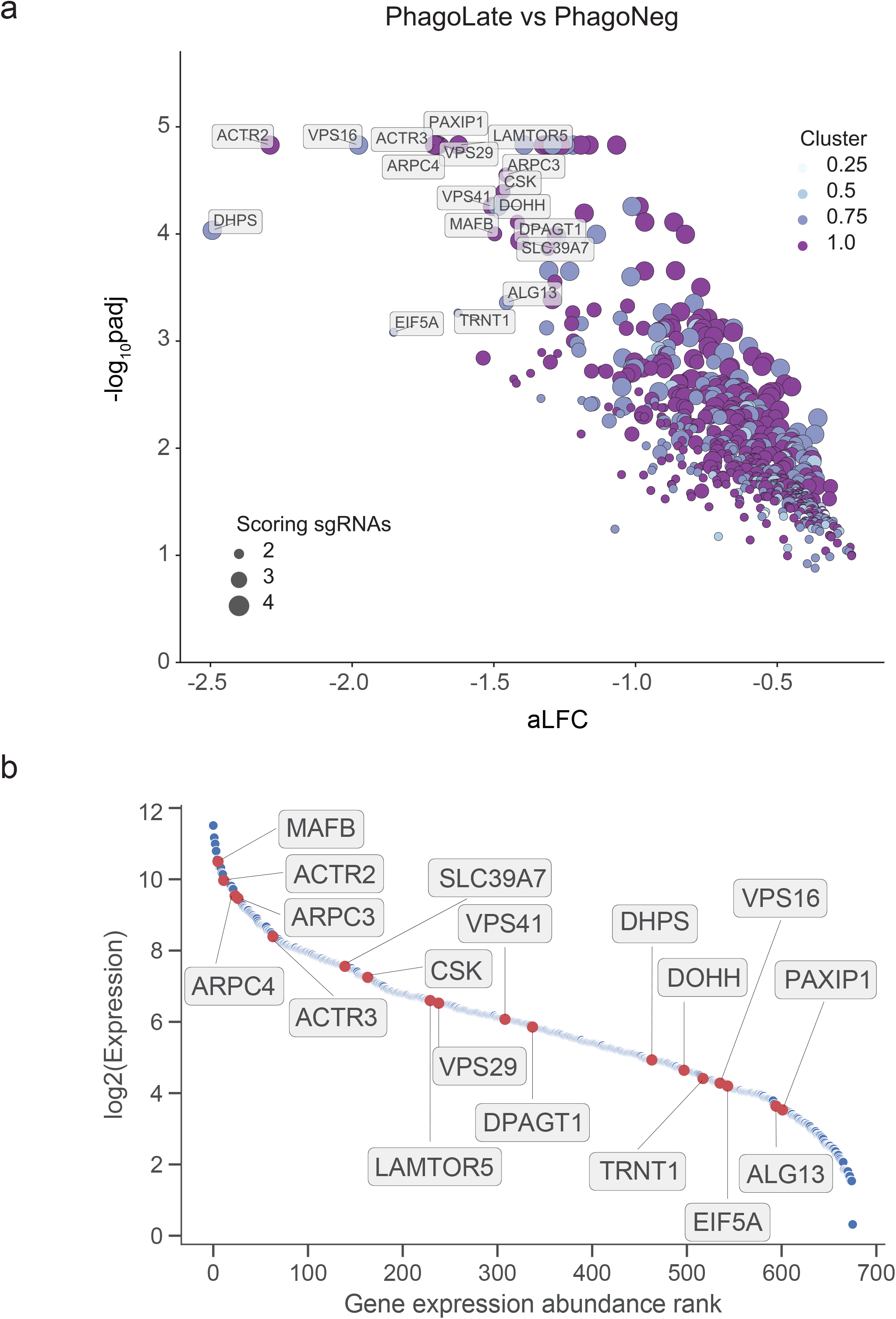
PhagoLate vs PhagoNeg CRIPRko hits. **a**. Volcano plot showing 716 genes depleted in the PhagoLate population versus the PhagoNeg population. The x-axis shows the average log_2_ -fold change (aLFC) calculated for all sgRNAs per gene against the y-axis, representing the statistical significance as -log_10_(padj). The size of the dot indicates the amount of sgRNAs changing significantly for the particular gene. The color indicates the expression level of the individual gene as measured by RNA sequencing of THP-1 cells stimulated with beads. Normalized expression levels of genes were categorized into the 0.25, 0.5, 0.75 and 1.0 quantiles and colored accordingly. **b**. Scatter plot showing the expression levels of all the 716 genes depleted in the PhagoLate population versus the PhagoNeg population as expressed in THP1 cells stimulated with beads. The x-axis shows the position of the gene ranked by ascending expression levels against the y-axis, representing the log_2_ transformed expression level of each gene. Highlighted in red are the same genes that are indicated in (a), representing the top downregulated genes of the screen.

**Figure S3:**
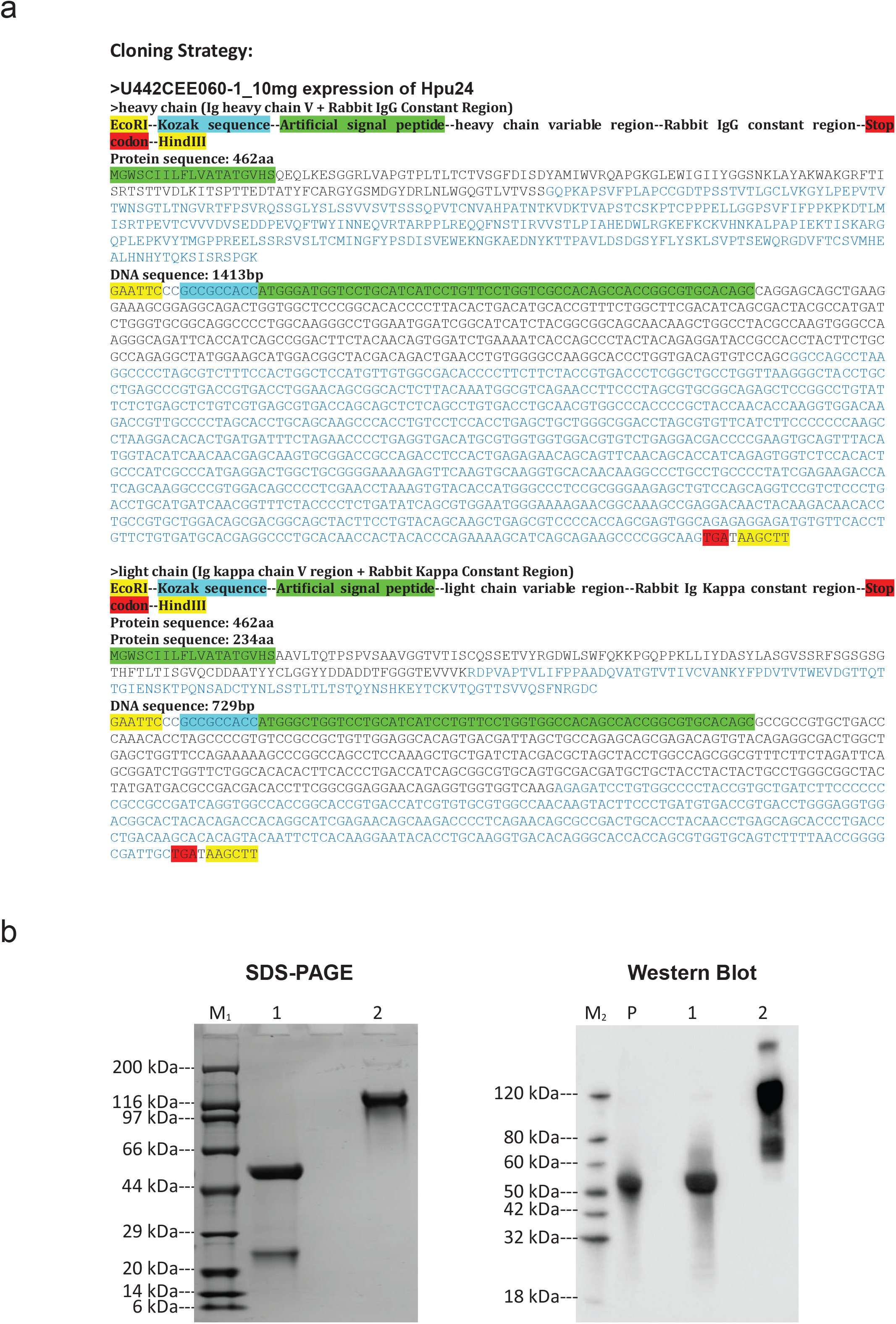
Generation of a hypusine binder. **a**. Cloning strategy for expression of the hypusine binder Hpu24. The coding sequences for the heavy and light chain of the antibody are shown as protein and DNA sequence. **b**. SDS-PAGE and Western blot controls. Analysis of purified antibody Hpu24 under reducing and non-reducing conditions. Western blot showing the detection of Rabbit IgG in reducing (lane 1) and non-reducing (lane 2) conditions for the purified antibody Hpu24 as well as for an IgG isolated from rabbit serum as positive control (lane P).

